# Hyperaerated metalation calculator for *E. coli* strain JM109 (DE3) grown in LB media

**DOI:** 10.1101/2022.09.02.506343

**Authors:** Sophie E. Clough, Deenah Osman, Tessa R. Young, Nigel J. Robinson

## Abstract

We recently produced three web-based calculators that predict *in vivo* metal occupancies of proteins, based on the metal affinities of a protein of interest along with estimates of the availabilities of the labile buffered pools of metals inside a cell. Metal availabilities were calculated from the calibrated responses of DNA-binding, metal-sensing, transcriptional regulators. The availability of intracellular Fe(II) was estimated to be similar in *E. coli* grown under anaerobic conditions compared to cells grown aerobically in LB medium. The purpose of this article is to archive the background data that underpins the release of a new calculator for hyperaerated cells grown in flasks with baffles, with relatively low culture volumes plus high shaking speeds to give elevated oxygenation. The intracellular availability of Fe(II) calculated from the responses of the intracellular Fe(II) sensor Fur was estimated to be significantly lower in these hyperaerated cells than either of the previous values determined for anaerobic or aerobic cultures. The total number of atoms of Mn(II) per cell increased in hyperaerated cells albeit with only modest change in intracellular Mn(II) availability as estimated from the responses of the Mn(II) sensor MntR. Accurate determination of intracellular Ni(II) availability will require further calibration of the magnitude of the responses of the Ni(II) sensor NikR in hyperaerated cells to take account of the state of Fnr. The hyperaerated metalation calculator is made available online and as a spreadsheet, for use by others.

## Introduction

To create metalation calculators for *E. coli* cells grown aerobically, anaerobically and exposed to hydrogen peroxide, the expression of selected genes that are regulated by known DNA binding, metal sensing, transcriptional regulators were previously monitored by qPCR [1]. Boundary conditions describing the highest and lowest levels of transcript abundance for each regulated gene were established. Transcript abundance was related to intracellular metal availability taking advantage of the measured thermodynamic properties of the related metal-sensors of *Salmonella enterica* serovar *Typhimurium* [2]. Intracellular metal availabilities were included in on-line calculators that simulate which metals will bind to a protein of known metal affinities at the respective metal availabilities when metalation approximates to thermodynamic equilibrium (https://mib-nibb.webspace.durham.ac.uk/metalation-calculators/).

There is variation in the response of some genes that are regulated by the Fe(II) sensor Fur [3]. A sub-set of Fur-responsive promoters show enhanced binding of the transcriptional regulator under anaerobic growth conditions compared to aerobic conditions. Notably, this did not include the *fepD* promoter which had been monitored to determine intracellular metal availabilities for the generation of metalation calculators [3]. Moreover, we speculated that it might be possible to observe oxygen-dependent differences in the expression of Fur-responsive promoters if cells were rigorously hyperaerated: Using flasks with internal baffles, small culture volumes to maximise the air interface, and high shaking speeds. Here we show that under these conditions *fepD* transcript abundance declines reflecting a substantial reduction in intracellular Fe(II) availability relative to cells grown anaerobically or aerobically but with less mixing. The availabilities of other metals have also been estimated within these hyperaerated cells allowing a further metalation calculator to be created, both as a spreadsheet provided here within the supplementary materials, and as an online resource (https://mib-nibb.webspace.durham.ac.uk/metalation-calculators/). This manuscript archives the data used to create the hyperaerated calculator.

## Methods

### Bacterial strain maintenance/growth and reagents

*E. coli* strain JM109 (DE3) was purchased from Promega. Liquid growth media and cultures were prepared in acid washed glassware or sterile plasticware to minimise metal contamination. Overnight cultures were inoculated into 400 ml LB (10 g/l tryptone, 5 g/l yeast extract, 10 g/l NaCl) at a 1 in 100 dilution and grown at 37 °C to an OD_600 nm_ of 0.2-0.3 then split into 10 ml aliquots in 100 ml acid washed baffled flasks with the indicated treatment and grown as indicated. The hyper-aerated cultures were incubated with shaking at 200 rpm for 2 h. OD_600 nm_ measurements were made using a Thermo Scientific Multiskan GO spectrophotometer with n = 3 biological replicates.

### Determination of transcript abundance

Aliquots (1 ml) of culture were added to RNAProtect Bacteria Reagent (Qiagen) (2 ml), RNA isolated a described previously [1], and cDNA generated using the SuperScript IV Reverse Transcriptase System (Invitrogen) with control reactions lacking reverse transcriptase prepared in parallel.

Transcript abundance was determined using primers 1 and 2 for *mntS*, 3 and 4 for *fepD*, 5 and 6 for *rcnA*, 7 and 8 for *nikA*, 9 and 10 for *znuA*, 11 and 12 for *zntA*, 13 and 14 for *copA*, 15 and 16 for *rpoD* designed to amplify ∼100 bp (Supplementary Table 1). qPCR was performed in 20 μl reactions containing 5 ng of cDNA, 400 nM of each primer and PowerUP SYBR Green Master Mix (Thermo Fisher Scientific). Three technical replicates of each biological sample were analysed as previously [1]. *C*_q_ values were calculated with LinReg PCR (version 2021.1). Samples were rejected where the *C*_q_ value for no template or –RT control was <10 (to the nearest integer) greater than the equivalent value for cDNA containing sample. These samples were either rerun or DNaseI treatment and cDNA synthesis performed again on RNA samples before running qPCR.

### Determination of boundary conditions for the expression of each transcript

Boundary conditions for the calibration of sensor response curves were defined by the minimum and maximum abundance of the regulated transcript.

Following completion of all qPCR reactions qPCR data for the control gene (*rpoD*) in each sample was reanalysed in a single LinReg PCR analysis along with collated data for *rpoD* expression in control conditions. The difference between *rpoD C*_q_ in the condition of interest and control condition was determined (average *rpoD C*_q_ for control condition minus average *rpoD C*_q_ for treated sample) (Supplementary Table 2). All hyperaerated samples were compared against previously generated untreated aerobic 2 h controls [1]. Notably a mean difference of 2.9 was found for *rpoD C*_q_ values, with a standard deviation of 0.1 most likely reflecting globally enhanced levels of gene expression in hyperaerated cells relative to previous aerobic cultures (Supplementary Table 3).

### Intracellular metal availability in conditional cells

The fold change in transcript abundance, relative to the mean of the control condition (lowest expression) for each sensor, was calculated as before [1]. Fractional responses of sensors were related to available metal concentrations via sensor response curves generated as before.

Intracellular available Δ*G*_Metal_ was calculated using equation 3 where [Metal] is the intracellular available metal concentration, *R* (gas constant) = 8.314 × 10^−3^ kJ K^-1^ mol^-1^ and *T* (temperature) = 298.15 K:

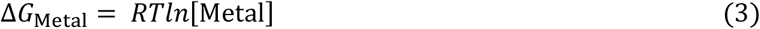

### Simulated metalation of molecules under bespoke conditions

A calculator has been created for prediction of molecule metalation in JM109 (DE3) under hyperaerated conditions using the same approaches as described previously (https://mib-nibb.webspace.durham.ac.uk/metalation-calculators/ and Supplementary Spreadsheet 1) [1].

### Determination of cellular metal quotas

Cells were grown in hyper-aerated conditions with a 2 h exposure to the indicated treatment before harvest (n = 3 biological replicates per treatment). Cell pellets were washed with 10 mM HEPES pH 7.8, 0.5 M sorbital, 1 mM EDTA followed by the same buffer without EDTA and digested in 65% HNO_3_ before analysis by ICP-MS. OD_600 nm_ values at harvest were converted to cells per ml using the approximation OD_600 nm_ = 1 correlates to 4.4 × 10^8^ cell ml^-1^ [4].

## Results

Figure 1 shows the fold change in *mntS, fepD, rcnA, nikA, znuA, zntA* and *copA* transcript abundance relative to the level of expression defined previously as the lower boundary condition for the respective promoter [1]. Data are also shown for maximum expression defined previously as the upper boundary condition (Supplementary Tables 4 – 10). Values for hyperaerated cells were within the two boundary conditions for all transcripts except *copA* where hyperaerated cells now define the lowest boundary: The average ΔCq value for *copA* transcripts in hyperaerated cells was 4.5 (+/- 0.1) while that previously reported for cells exposed to H_2_O_2_ was 3.4 (+/-0.3), the latter having been used to define the lower boundary condition. Future experiments could coincidentally compare *copA* transcript abundance in hyperaerated versus H_2_O_2_ exposed cells, with the formal possibility that estimates of intracellular Cu(I) availabilities in aerobic, anaerobic and H_2_O_2_ exposed cultures might slightly increase. The fold changes in gene expression shown in Figure 1 were calibrated to DNA occupancies (conditional *θ*_D_ for repressors or *θ*_DM_ for activators) and DNA occupancies related to buffered metal concentration as described previously [1], to generate Figure 2.

**Figure 1.**
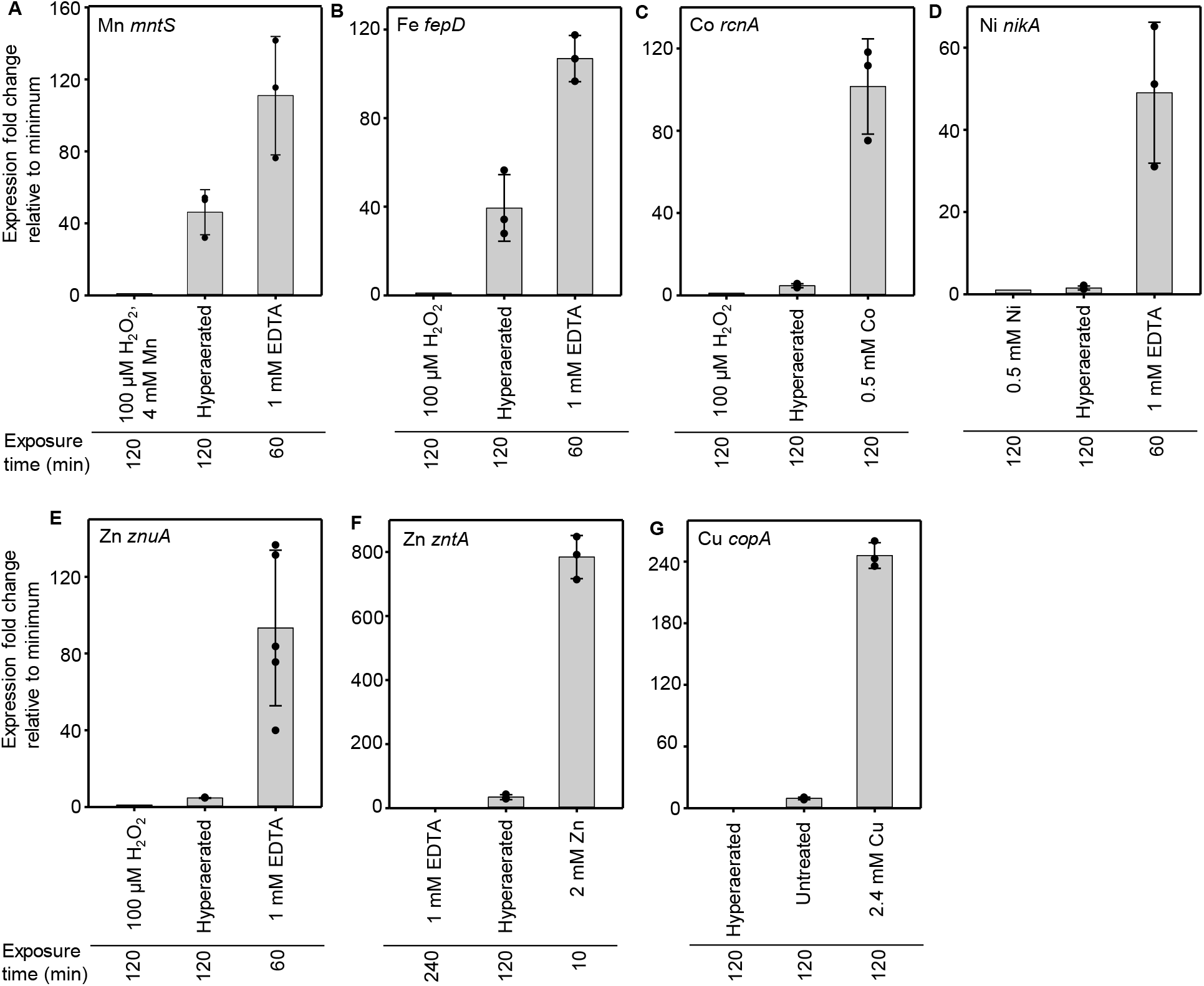
Fold change in metal responsive transcript abundance under hyper-aerated conditions relative to a minimum. Abundance of seven metal-responsive transcripts grown under hyper-aerated conditions relative to the first (left) condition shown on each chart (assigned a value of 1) as determined by qPCR. Note that in panel G the hyper-aerated transcript abundance was the lowest and assigned the value of 1. Also note the larger scales in panels F and G. A. *mntS* regulated by Mn^2+^ sensor MntR. B. *fepD* regulated by Fe^2+^ sensor Fur. C. *rcnA* regulated by Co^2+^ sensor RcnR. D. *nikA* regulated by Ni^2+^ sensor NikR (aerobic growth). E. *znuA* regulated by Zn^2+^ sensor Zur. F. *zntA* regulated by Zn^2+^ sensor ZntR. G. *copA* regulated by Cu^+^ sensor CueR.

**Figure 2.**
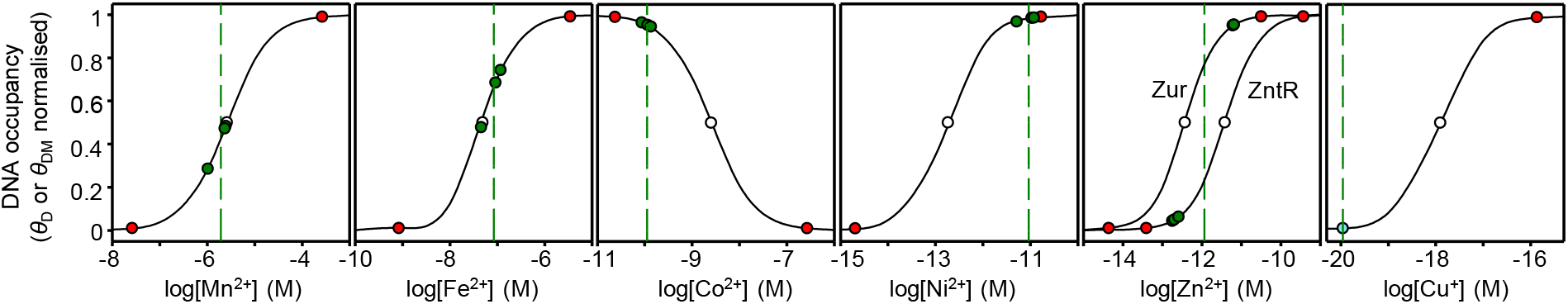
Calibrated responses of metal sensors as a function of intracellular metal-availability under hyper-aerated conditions. The calculated relationship between intracellular metal availability and normalised DNA-occupancy (*θ*_D_), or for activators normalised DNA-occupancy with metalated sensor (*θ*_DM_). The dynamic range within which each sensor responds to changing intracellular metal availability has been defined as *θ*_D_ or *θ*_DM_ of 0.01 to 0.99 representing the boundary conditions for maximum or minimum fold-changes in transcript abundance (red circles). Mid-point of each range (open circle), replicate values calculated for LB media under hyper-aerated conditions (green circles), and mean availability (green dashed lines). For Zn^2+^ two curves represent the responses of *znuA* and *zntA*, here the green dashed line is a midpoint between the means for each sensor. Note that for Cu^+^ the minimum boundary condition was defined by the hyper-aerated value (blue circle). Buffered concentrations and curves were calculated as described by Foster and co-workers [1].

Buffered concentrations of available metals inside hyperaerated cells (indicated by the dashed green lines on Figure 2) were numerically derived from the previously established relationships to *θ*_D_ or *θ*_DM_ for each sensor using MATLAB code (Supplementary Note 3) available in Osman and co-workers [2], and the values are shown in Table 1. Ni(II) is an exception in that expression from the *nikA* promoter is also subject to control by Fnr. A decision was made to populate Table 1 with the previously determined Ni(II) availability in aerobically cultured cells as an interim measure, noting that high and low boundary conditions would need to be independently calibrated for hyperaerated cells subjected to Ni(II) supplementation, and separately to depletion using a chelator.

**Table 1.** Calculated available metal concentrations in hyperaerated cells^a^.

| Metal | Sensor | Hyper-aerated<br>[Metal]<sub>buffered</sub> (M) |
| --- | --- | --- |
| Mn<sup>2+</sup> | MntR | 1.9(±0.8) × 10<sup>-6</sup> |
| Fe<sup>2+</sup> | Fur | 8.4(±4) × 10<sup>-8</sup> |
| Co<sup>2+</sup> | RcnR | 1.1(±0.2) × 10<sup>-10</sup> |
| Zn<sup>2+</sup> | Zur | 6.3(±0.2) × 10<sup>-12</sup> |
|  | ZntR | 2.2(±0.4) × 10<sup>-13</sup> |
|  | Mid-point | 1.2 × 10<sup>-12</sup> |
| Ni<sup>2+</sup> | NikR | 1.3(±0.7) × 10<sup>-13</sup> |
| Cu<sup>+</sup> | CueR | 1.1 × 10<sup>-20</sup> |

<sup>a</sup> Values were initially calculated to 2 significant figures in supplementary spreadsheets and calculators. Here the finalised numbers have been rounded and the number of decimal places rationalised to those of the standard deviation (shown in parenthesis).
<sup>b</sup> Boundary conditions do not have standard deviations

Figure 3 shows metal availabilities in hyperaerated cells as free energies for forming complexes with proteins (or other types of molecules) that will be 50% metalated at the respective buffered available metal concentrations shown in Table 1. Standard deviations were calculated based upon the triplicated determinations of buffered concentration which have been averaged in Table 1. For reasons described above, Ni(II) in hyperaerated cells has been excluded from Figure 3. For comparison, previously used metal-availabilities at the mid-points of the ranges of each sensor for each metal in idealised cells, are also shown (grey symbols in Fig. 3). For comparison with intracellular metal availabilities, Figure 4 shows the total number of atoms per cell of Mn, Fe, Ni, Cu, Zn (Co was below the calibrated range for detection) in cultures grown under hyperaerated conditions and under the more standard aerobic conditions that had been used previously.

**Figure 3.**
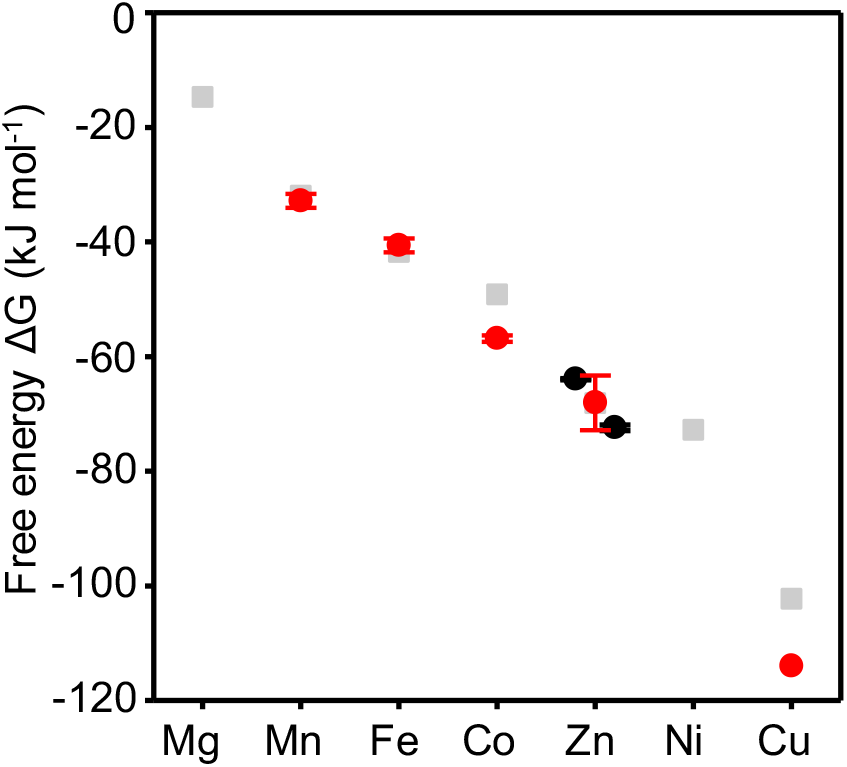
Free energies of available metal in *E. coli* JM109 (DE3) under hyper-aerated conditions. Intracellular available free energies for metal binding to molecules that would be 50 % saturated at the available metal concentration (Table 1) in hyperaerated cells (red circles with standard deviation). For Zn^2+^ separate values have been derived based on the response of Zur and ZntR (black circles with standard deviation, Zur = left, ZntR = right). For Ni^+^ the red circle has not been represented as the boundary conditions would need to be redefined. Grey squares are values for idealised cells reflecting the mid-point of sensor ranges as used in previous versions of calculators [4].

**Figure 4.**
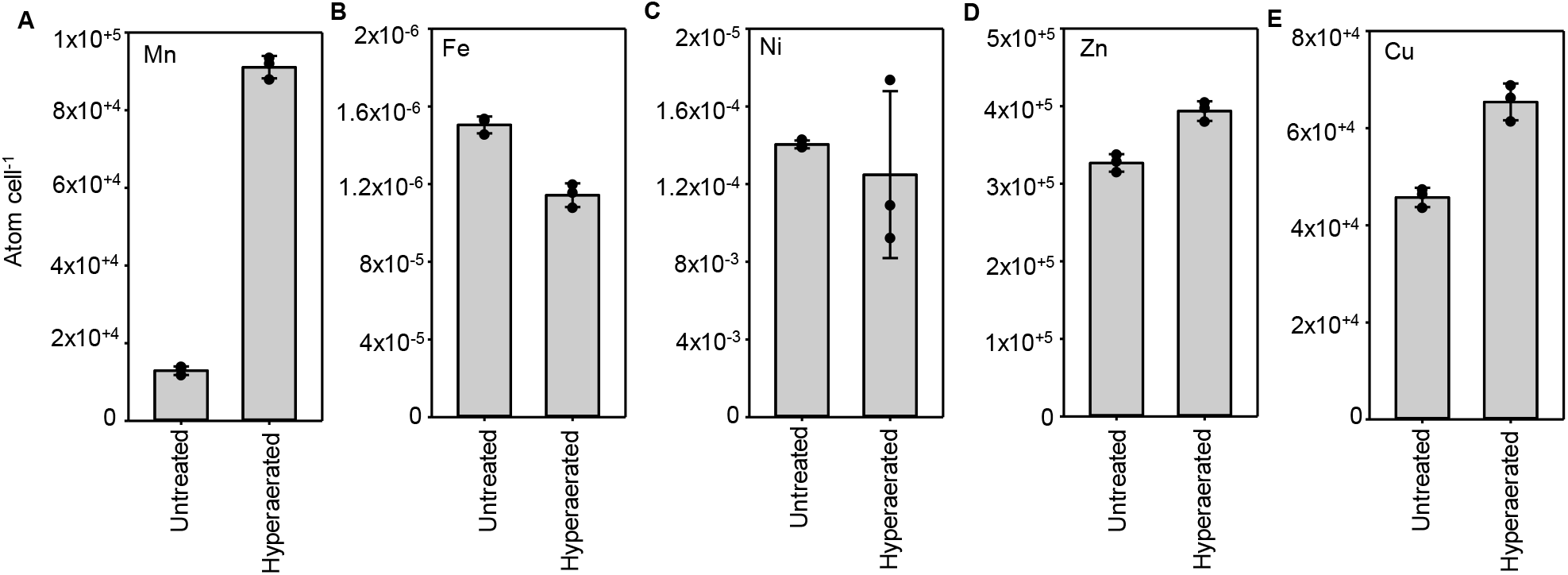
Total cellular metal content of *E. coli* JM109(DE3) under hyper-aerated conditions. Cells were cultured in hyper-aerated conditions with a 2 h exposure to the indicated treatment before harvest (n = 3 biological replicates per treatment). Data is presented as mean ± standard deviation. Note the magnitude of scale difference between **A-E**.

To predict the metalation states of proteins of known metal affinities, a metalation calculator was created based on the intracellular metal availabilities estimated in hyperaerated cells (Figure 3 and Table 1). The calculator was produced as a spreadsheet (Supplementary Spreadsheet 1), and as a web-based version (https://mib-nibb.webspace.durham.ac.uk/metalation-calculators/), using previously described approaches [1]. Metals (for example Ni(II)) can be excluded from the calculations where there is uncertainty and enabling simulations for proteins where some affinities are unknown. Metal affinities of proteins of interest are entered as dissociation constants, *K*_D_.

## Discussion

Metal availability in hyperaerated cells, in common with idealised cells, follows the Irving Williams series (Figure 3, Table 1). The availabilities of Co(II) and Cu(I) are lower than in idealised cells and a similar observation was previously reported in cells grown anaerobically, aerobically or exposed to H_2_O_2_ [1]. Most importantly, in cells grown anaerobically, aerobically or exposed to H_2_O_2_, the availability of Fe(II) was greater than idealised cells, whereas hyperaerated cells contain a significantly reduced intracellular Fe(II) availability, more similar to that of idealised cells.

Intracellular Mn(II) is slightly less available in hyperaerated cells compared to previous estimates in aerated cultures (Figure 3, Table 1) [1]. However, the number of Mn(II) atoms per cell greatly increase under hyperaerated conditions (Figure 4). It is anticipated that reactive oxygen species increase in hyperaerated cells triggering increased expression of Mn(II) import via the known actions of the OxyR sensor [5]. Mn(II) is a potent antioxidant and required by the SodA Mn(II)-dependent superoxide dismutase. Intriguingly, these data suggest that the demand for Mn(II) slightly depletes the buffered bioavailable level even though accumulation in Mn(II) proteins leads to a corresponding increase in the overall cellular Mn(II) quota. Moreover, the decline in available Fe(II) within hyperaerated cells is suggested to be linked to the increase in total Mn(II) and a response to reactive oxygen species.

## Acknowledgements

This work was supported by Biotechnology and Biological Sciences Research Council awards: BB/S009787/1, BB/V006002/1, BB/W015749/1 a Royal Commission for Exhibition of 1851 Research Fellowship (TRY).

## Data availability

The data underlying this article are available in the article and in its online supplementary material.

