## Supplementary information for "Hyperaerated metalation calculator for *E. coli* strain JM109 (DE3) grown in LB media"

**Supplementary Table 1**. qPCR primers used in this work.

| No. | Name | Sequence | Product size (bp) | Reference |
|---|---|---|---|---|
| 1 | *mntS*_F | 5'-GTATGCGCGTGTTTAGTCATTC-3' | 105 | Ref [1] |
| 2 | *mntS*_R | 5'-TATCGGAAGGTTTATCTTGCTG-3' | 105 | Ref [1] |
| 3 | *fepD*_F | 5'-TGCAAACCCTCACCCGAAAC-3' | 111 | Ref [1] |
| 4 | *fepD*_R | 5'-GCGCGGAAGAGTAACCAAACAG-3' | 111 | Ref [1] |
| 5 | *rcnA*_F | 5'-GAACCAGGGCACTCAAAAAC-3' | 108 | Ref [2] |
| 6 | *rcnA*_R | 5'-TGCGGTATGCGAAATAGTTG-3' | 108 | Ref [2] |
| 7 | *nikA*_F | 5'-AACCCGCACCTTTACACGCC-3' | 114 | Ref [1] |
| 8 | *nikA*_R | 5'-AGTCCAGCTTTTTGCCAGCC-3' | 114 | Ref [1] |
| 9 | *znuA*_F | 5'-GTTTGGACTGACACCGCTTG-3' | 111 | Ref [3] |
| 10 | *znuA*_R | 5'-ACGCAGGTTGCTTTTTGCTC-3' | 111 | Ref [3] |
| 11 | *zntA*_F | 5'-CGAAGCACAGGTTGCTGAAC-3' | 114 | Ref [1] |
| 12 | *zntA*_R | 5'-CCGGCAGCGCAAATCAATAC-3' | 114 | Ref [1] |
| 13 | *copA*_F | 5'-GTCACAAACTATCGACCTGACCC-3' | 104 | Ref [1] |
| 14 | *copA*_R | 5'-CATCCGCCTGCTCAACATCC-3' | 104 | Ref [1] |
| 15 | *rpoD*_F | 5'-GTGGCTTGCAGTTCCTTGAC-3' | 108 | Ref [3] |
| 16 | *rpoD*_R | 5'-AGGTTGCGTAGGTGGAGAAC-3' | 108 | Ref [3] |

**Supplementary Table 2**. $\Delta C_q$ values obtained from qPCR analysis of hyperaerated samples with all target primers and *rpoD* (control)[a].

| Target primer | Exposure time (min) | $\Delta C_{q\ target\text{-}rpoD}$ |
|---|---|---|
| *mntS* | 120 | 1.4(±0.4) |
| *fepD* | 120 | 3.3(±0.5) |
| *rcnA* | 120 | 7.3(±0.3) |
| *nikA* | 120 | 6.6(±0.4) |
| *znuA* | 120 | 2.1(±0.05) |
| *zntA* | 120 | 2.2(±0.3) |
| *copA* | 120 | 4.5(±0.1) |

[a] $\Delta C_{q\ target\text{-}rpoD}$ obtained for each biological replicate, mean (± standard deviation).

**Supplementary Table 3**. *rpoD* $C_q$ difference values[a].

| Treatment | Exposure time (min) | *rpoD* $C_q$ difference relative to untreated aerobic samples |
|---|---|---|
| Hyperaerated | 120 | 2.93(±0.1) |

[a] *rpoD* $C_q$ difference between values determined for hyperaerated samples and the mean value determined for untreated aerobic cultures. Values shown as mean ± standard deviation.

**Supplementary Table 4**. $\Delta\Delta C_q$ values for conditional response of *mntS* promoter[a].

| $O_2$ | Treatment | Exposure time (min) | $\Delta\Delta C_q$ |
|---|---|---|---|
| $+O_2$ | 1 mM EDTA | 60 | -6.7(±0.5) |
| $++O_2$ | Hyperaerated | 120 | -5.5(±0.4) |
| $+O_2$ | 100 µM $H_2O_2$, 4 mM Mn | 60 | - |

[a] Determined by subtraction of $\Delta C_q$ for control condition (lowest expression, aerobic 60 min exposure to 100 µM $H_2O_2$ and 4 mM Mn, Supplementary Table 8 in [1]) from $\Delta C_q$ for the condition of interest.

**Supplementary Table 5**. $\Delta\Delta C_q$ values for conditional response of *fepD* promoter[a].

| $O_2$ | Treatment | Exposure time (min) | $\Delta\Delta C_q$ |
|---|---|---|---|
| $+O_2$ | 1 mM EDTA | 60 | -6.7(±0.1) |
| $++O_2$ | Hyperaerated | 120 | -5.2(±0.5) |
| $+O_2$ | 100 µM $H_2O_2$ | 120 | - |

[a] Determined by subtraction of $\Delta C_q$ for control condition (lowest expression, aerobic 120 min exposure to 100 µM $H_2O_2$, Supplementary Table 9 in [1]) from $\Delta C_q$ for the condition of interest.

**Supplementary Table 6**. $\Delta\Delta C_q$ values for conditional response of *rcnA* promoter[a].

| $O_2$ | Treatment | Exposure time (min) | $\Delta\Delta C_q$ |
|---|---|---|---|
| $+O_2$ | 0.5 mM Co | 120 | -6.6(±0.4) |
| $++O_2$ | Hyperaerated | 120 | -3.7(±0.2) |
| $+O_2$ | 100 µM $H_2O_2$ | 120 | - |

[a] Determined by subtraction of $\Delta C_q$ for control condition (lowest expression, aerobic 120 min exposure to 100 µM $H_2O_2$, Supplementary Table 10 in [1]) from $\Delta C_q$ for the condition of interest.

**Supplementary Table 7**. $\Delta\Delta C_q$ values for conditional response of *nikA* promoter under aerobic conditions[a].

| $O_2$ | Treatment | Exposure time (min) | $\Delta\Delta C_q$ |
|---|---|---|---|
| $+O_2$ | 1 mM EDTA | 60 | -5.6(±0.5) |
| $++O_2$ | Hyperaerated | 120 | -2.1(±0.2) |
| $+O_2$ | 0.5 mM Ni | 120 | - |

[a] Determined by subtraction of $\Delta C_q$ for control condition (lowest expression, aerobic 120 min exposure to 0.5 mM Ni, Supplementary Table 7 in [1]) from $\Delta C_q$ for the condition of interest.

**Supplementary Table 8**. $\Delta\Delta C_q$ values for conditional response of *znuA* promoter[a].

| $O_2$ | Treatment | Exposure time (min) | $\Delta\Delta C_q$ |
|---|---|---|---|
| $+O_2$ | 1 mM EDTA | 60 | -6.4(±0.7) |
| $++O_2$ | Hyperaerated | 120 | -2.2(±0.04) |
| $+O_2$ | 100 µM $H_2O_2$ | 120 | - |

[a] Determined by subtraction of $\Delta C_q$ for control condition (lowest expression, aerobic 120 min exposure to 100 µM $H_2O_2$, Supplementary Table 11 in [1]) from $\Delta C_q$ for the condition of interest.

**Supplementary Table 9**. $\Delta\Delta C_q$ values for conditional response of *zntA* promoter[a].

| $O_2$ | Treatment | Exposure time (min) | $\Delta\Delta C_q$ |
|---|---|---|---|
| $+O_2$ | 2 mM Zn | 10 | -9.6(±0.1) |
| $++O_2$ | Hyperaerated | 120 | -5.1(±0.3) |
| $+O_2$ | 1 mM EDTA | 240 | - |

[a] Determined by subtraction of $\Delta C_q$ for control condition (lowest expression, aerobic 240 min exposure to 1 mM EDTA, Supplementary Table 12 in [1]) from $\Delta C_q$ for the condition of interest.

**Supplementary Table 10**. $\Delta\Delta C_q$ values for conditional response of *copA* promoter[a].

| $O_2$ | Treatment | Exposure time (min) | $\Delta\Delta C_q$ |
|---|---|---|---|
| $+O_2$ | 2.4 mM Cu | 120 | -6.8(±0.07) |
| $++O_2$ | Hyperaerated | 120 | - |

[a] Determined by subtraction of $\Delta C_q$ for control condition (lowest expression, Supplementary Table 13 in [1]) from $\Delta C_q$ for the condition of interest.
